# Coronavirus genomes carry the signatures of their habitats

**DOI:** 10.1101/2020.06.13.149591

**Authors:** Yulong Wei, Jordan R. Silke, Parisa Aris, Xuhua Xia

## Abstract

Coronaviruses such as SARS-CoV-2 regularly infect host tissues that express antiviral proteins (AVPs) in abundance. Understanding how they evolve to adapt or evade host immune responses is important in the effort to control the spread of COVID-19. Two AVPs that may shape viral genomes are the zinc finger antiviral protein (ZAP) and the apolipoprotein B mRNA-editing enzyme-catalytic polypeptide-like 3 protein (APOBEC3). The former binds to CpG dinucleotides to facilitate the degradation of viral transcripts while the latter deaminates C into U residues leading to dysfunctional transcripts. We tested the hypothesis that both APOBEC3 and ZAP may act as primary selective pressures that shape the genome of an infecting coronavirus by considering a comprehensive number of publicly available genomes for seven coronaviruses (SARS-CoV-2, SARS-CoV, MERS, Bovine CoV, Murine MHV, Porcine HEV, and Canine CoV). We show that coronaviruses that regularly infect tissues with abundant AVPs have CpG-deficient and U-rich genomes; whereas viruses that do not infect tissues with abundant AVPs do not share these sequence hallmarks. In SARS-CoV-2, CpG is most deficient in the S protein region to evaded ZAP-mediated antiviral defense during cell entry. Furthermore, over four months of SARS-CoV-2 evolutionary history, we observed a marked increase in C to U substitutions in the 5’ UTR and ORF1ab regions. This suggests that the two regions could be under constant C to U deamination by APOBEC3. The evolutionary pressures exerted by host immune systems onto viral genomes may motivate novel strategies for SARS-CoV-2 vaccine development.

## INTRODUCTION

The emergence of SARS-CoV-2 pandemic poses a serious global health emergency. understanding how coronaviruses adapt or evade tissue-specific host immune responses is, therefore, important in the effort to control the spread of COVID-19 and to facilitate vaccine-development strategies. As obligate parasites, coronaviruses evolve in mammalian hosts and carry genomic signatures shaped by their host-specific environments. At the tissue level, mammalian species provide different cellular environments with varying levels of antiviral and RNA modification activity. Two antiviral proteins (AVPs) that may contribute to the modification of viral genomes are the zinc finger antiviral protein (ZAP, gene name ZC3HAV1 in mammals) and the apolipoprotein B mRNA-editing enzyme-catalytic polypeptide-like 3 (APOBEC3), both of which exhibit tissue-specific expression (Fagerberg *et al.* 2014).

ZAP is a key component in the mammalian interferon-mediated immune response that specifically targets CpG dinucleotides in viral RNA genomes (Meagher *et al.* 2019) to inhibit viral replication and signal for viral genome degradation (Ficarelli *et al.* 2020; Guo *et al.* 2007; Meagher *et al.* 2019; Takata *et al.* 2017). This host immune response acts against not only retroviruses such as HIV-1 (Ficarelli *et al.* 2020; Zhu *et al.* 2011), but also single-stranded RNA viruses such as Ecovirus 7 (Odon *et al.* 2019), Zika virus (Trus *et al.* 2020), and Influenza virus (Greenbaum *et al.* 2008). It follows that cytoplasmic ZAP activity should impose a strong avoidance of CpG dinucleotides in RNA viruses that target host tissues abundant in ZAP. For instance, while HIV-1 infects lymph organs where ZAP is abundant (Fagerberg *et al.* 2014), its genome is also strongly CpG-deficient. Notably, the viral fitness of HIV-1 has been shown to diminish as its genomic CpG content increases within a sample of patients (Theys *et al.* 2018). Many other pathogenic single-stranded RNA viruses, including coronaviruses, also exhibit strong CpG deficiency (Atkinson *et al.* 2014; Greenbaum *et al.* 2008; Greenbaum *et al.* 2009; Takata *et al.* 2017; Yap *et al.* 2003), but selection for CpG deficiency disappears in ZAP-deficient cells (Takata *et al.* 2017).

The ZAP-mediated RNA degradation is cumulative (Takata *et al.* 2017). When CpG dinucleotides are added to individual viral segment 1 or 2 in HIV-1, the inhibitory effect of ZAP is weak. However, when the same CpG dinucleotides are added to both segments 1 and 2, the ZAP inhibition effect is strong (Takata *et al.* 2017). This implies that only RNA sequences of sufficient length would be targeted by ZAP. Thus, although SARS-CoV-2 has the lowest genomic CpG contents found in Betacoronaviruses (Xia 2020), only ORF1aband spike (S) mRNAs are likely targets by ZAP. It is therefore not surprising that these mRNAs, especially the S mRNA, exhibit lower CpG than other shorter genes (di Gioacchino *et al.* 2020; Kim *et al.* 2020).

Aside from ZAP, the APOBEC3 cytidine deaminase enzymes have garnered substantial attention for their role in the antiviral immune response (Cullen 2006; Harris and Dudley 2015). Through a mechanism largely derived from HIV-1 studies, APOBEC3 enzymes have been prominently reported to disrupt the structure and function of HIV-1 viruses by hypermutating minus strand viral cDNA, causing defects in the viral transcript and inhibiting reverse transcription (Chiu and Greene 2008; Harris and Dudley 2015; Hayward *et al.* 2018; Nabel *et al.* 2013; Rodriguez-Frias *et al.* 2013). For instance, APOBEC3G (Harris *et al.* 2003; Mangeat *et al.* 2003) and 3F (Zheng *et al.* 2004) catalyzes C to U deamination at the HIV-1 minus-strand DNA during reverse transcription, this triggers G to A hypermutation in the nascent retroviral DNA. HIV-1 avoids these deleterious effects by expressing Vif, a protein which targets and degrades APOBEC3 enzymes (Sheehy *et al.* 2002; Wang and Wang 2009). Despite these findings, the possibility that APOBEC3 paralogues may act directly to edit ssRNA viruses has not been widely explored but the potential should not be excluded, especially for viruses that do not encode a Vif analogue such as SARS-CoV-2.

Indeed, APOBEC3 is now known to modify a variety of RNA sequences. For instance, RNA binding activity facilitates the packaging of APOBEC3 into virions (Bogerd and Cullen 2008; Zhang *et al.* 2010). Furthermore, C to U RNA editing has been demonstrated in macrophages, monocytes, and lymphocytes by both APOBEC3A and 3G (Sharma *et al.* 2016; Sharma *et al.* 2015; Sharma *et al.* 2019). Additionally, APOBEC3C, 3F, and 3H may inhibit HCoV-NL63 coronavirus infection in humans, yet it remains unclear whether C to U RNA editing was involved (Milewska *et al.* 2018). More recently, studies have shown evidence that SARS-CoV-2 genomes are driven towards increasing U content and decreasing C content (Di Giorgio *et al.* 2020; Jiang 2020; Simmonds 2020; Victorovich *et al.* 2020). Resultantly, the possibility of C-U editing by APOBEC3 at the RNA level could effectively disrupt the structure and protein function of positive single-stranded RNA viruses.

We hypothesize that both APOBEC3 and ZAP act in concert as the primary selective pressure driving the adaptation of an infecting coronavirus over the course of its evolutionary history in specific host tissues. To test this hypothesis, we examined which antiviral proteins are effective against coronaviruses and how the immune response is subverted. We predict that when a virus regularly infects host tissues that are deficient in AVPs, there will be no strong directional substitutions resulting in decreased CpG dinucleotides or elevated U residues, as these evolutionary forces will be weak when ZAP and APOBEC3 are lowly expressed. Conversely, when a virus regularly infects host tissues that are abundant in AVPs, these antiviral responses will exert their influence on viral genomes. Consequently, viral genomes should tend towards reduced CpG dinucleotides to elude ZAP-mediated cellular antiviral defense, and increased U residues because of RNA editing by APOBEC3 proteins.

Our investigation considers a comprehensive number of publicly available genomes for seven coronaviruses (the Betacoronaviruses SARS-CoV-2, SARS-CoV, MERS, Bovine CoV, Murine MHV, and Porcine HEV, and the Alphacoronavirus Canine CoV,) as well as studies with tissue-specific ZAP and APOBEC3 gene expressions in five host species (human, cattle, dog, mice, and pig). We found that all surveyed coronaviruses regularly infect tissues with high mRNA expressions of both ZAP and APOBEC3, except Murine MHV. Expectedly, all surveyed coronaviruses, except Murine MHV, have high global CpG deficiency, with SARS-CoV-2 genomes having the lowest CpG content. More specifically, we observed a nonuniform distribution of CpG content across 12 SARS-CoV-2 viral regions and noted that CpG is most deficient in the region encoding the S protein that mediates cell entry by ACE2 binding. Taken together, these observations suggest SARS-CoV-2 has evolved in a tissue with high ZAP expression, and its persistence indicates that it has successfully evaded the ZAP-mediated antiviral defense during cell entry.

In line with evidence of RNA-level C to U deamination by APOBEC3 enzymes (Bishop *et al.* 2004; Sharma *et al.* 2016; Sharma *et al.* 2015; Sharma *et al.* 2019), Bovine CoV, Canine CoV, and Porcine HEV all exhibit high global U content and low global C content whereas the genomes of Murine MHV and the much more recent human coronaviruses (SARS-CoV-2, SARS-CoV, and MERS) exhibit notably lower U content and higher C content. To elucidate the early stages of SARS-CoV-2 genomic evolution, we analyzed the patterns of single nucleotide polymorphisms (SNPs) in local viral regions of complete genomes that were collected in the span of four months (from December 31, 2019 to May 6, 2020) since SARS-CoV-2 was initially isolated. We observed that the occurrence of C to U substitutions is strikingly more prevalent than any other SNPs, especially in the 5’ UTR and ORF1ab regions. This suggests that the 5’ UTR and ORF1ab regions are under constraint by these enzymes. Indeed, both APOBEC3 and ZAP exert selective pressure on the RNA genome compositions of coronaviruses that regularly infect tissues expressing the two antiviral genes in abundance, but they do not affect the RNA genomes of viruses that avoid infecting tissues with high antiviral gene expression.

## MATERIALS AND METHODS

### Retrieving and processing the *APOBEC3* and *ZAP* genes and their tissue specific gene expressions in five mammalian species

The NCBI Nucleotide Database was queried for all records containing “APOBEC3” and “ZC3HAV1L” as gene names, “Mammalia” as a taxonomic class, and *“Homo sapiens”, “Bos taurus”, “Canis lupus familiaris”, “Mus musculus”*, and *“Sus scrofa”* as species. These five species were selected because they have extensive tissue-specific gene expression studies (as discussed below). Next, each entry was searched for /product= ‘apolipoprotein B mRNA editing enzyme, catalytic polypeptide 3’ and for /product= ‘zinc finger CCCH-type containing, antiviral 1’, whole-genome and chromosome-wide results were excluded, and only the coding DNA sequence region of APOBEC3 and ZC3HAV1 isoforms were extracted in FASTA format along with their ENSEMBL Accession IDs.

To compare gene expressions of APOBEC3 and ZC3HAV1L among tissues, we retrieved publicly available RNA Sequencing and Microarray studies that each sampled at least 10 mammalian tissues. The five mammalian species that have extensive tissue-specific mRNA expressions are *Homo sapiens, Bos taurus, Canis lupus familiaris, Mus musculus*, and *Sus scrofa.* For *Homo sapiens*, tissue-specific mRNA expressions were retrieved in averaged FPKM values from all 171 RNA-Seq datasets in BioProject PRJEB4337(Fagerberg *et al.* 2014), 48 RNA-Seq datasets in BioProject PRJEB2445, 20 RNA-Seq datasets in BioProject PRJNA280600 (Duff *et al.* 2015), and in median TPM values from all RNA-Seq datasets available in the GTEx Portal (Lonsdale *et al.* 2013). For *Mus musculus*, tissue-specific mRNA expressions were retrieved in averaged FPKM values from all 741 RNA-Seq datasets in BioProject PRJNA66167 (mouse ENCODE consortium) (Yue *et al.* 2014) and in average TPM values from all 79 RNA-Seq datasets in BioProject PRJNA516470 (Naqvi *et al.* 2019). For *Sus scrofa*, tissue-specific mRNA expressions were retrieved in averaged FPKM values from TISSUE 2.0 integrated datasets (Palasca *et al.* 2018). For *Canis lupus familiaris*, tissue-specific gene expressions were retrieved in averaged fluorescence intensity units (FIU) from all 39 microarray datasets in BioProject PRJNA124245 (Briggs *et al.* 2011), and in averaged TPM values from all 75 RNA-Seq datasets in BioProject PRJNA516470 (Naqvi *et al.* 2019). Lastly, for *Bos taurus*, tissue-specific mRNA expressions were retrieved in averaged FPKM values from 42 RNA-Seq datasets in the Bovine Genome Database (Shamimuzzaman *et al.* 2019).

Given that the data extracted were from multiple independent sources, thus not directly comparable, the relative mRNA expression level designations (high or low) for APOBEC3 and ZAP isoforms in a given tissue were derived from comparisons among AVP expressions in all tissues in each independent source. Specifically, we calculated the proportion of mRNA expression (PME) as:

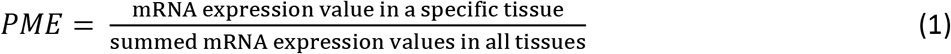

PME values were calculated from averaged TPM values in 24 human tissues using all RNA-Seq datasets available in the GTEx Portal (Lonsdale *et al.* 2013), from averaged FPKM values in 26 cattle tissues using the Bovine Genome Database (Shamimuzzaman *et al.* 2019), from averaged FPKM values in 33 pig tissues using TISSUE 2.0 integrated datasets (Palasca *et al.* 2018), from averaged FPKM values in 17 mice tissues using all 741 RNA-Seq datasets in mouse ENCODE consortium (Yue *et al.* 2014), from averaged FPKM values in 12 mice tissues using 79 RNA-Seq datasets in BioProject PRJNA516470 (Naqvi *et al.* 2019), and from averaged fluorescence intensity units in 10 dog tissues using all 39 microarray datasets in BioProject PRJNA124245 (Briggs *et al.* 2011). Next, we calculated the averaged PME value by considering all tissue-specific PME values in each independent source. Finally, for each AVP, tissue-specific PMEs were designated as high if they are greater than the averaged PME value and low if they are less than the averaged PME. In addition, each column in Supplemental figures S1 and S2 with column title designations “APOBEC3” or “ZC3HAV1” contains the tissue-specific AVP expressions from an individual source, where darkest blue represents the tissue with the highest mRNA expression and darkest red represents the lowest mRNA expression.

### Retrieving and processing the genomes and regular habitats of coronaviruses infecting five mammalian species

The genome, Accession ID, and Sample Collection Date of 28475 SARS-CoV-2 samples were retrieved from the China National Center for Bioinformation (CNCB) (https://bigd.big.ac.cn/ncov/variation/statistics?lang=en, last accessed May 16, 2020), among which 2666 strains were selected because they were annotated as having complete genome sequences and high sequencing quality. Additionally, the complete genomic sequences of 403 MERS strains, 134 SARS-CoV strains, 20 Bovine CoV strains, 2 Canine CoV strains, 26 Murine HEV strains, and 10 Porcine HEV strains were downloaded from the National Center for Biotechnology Information (NCBI) Nucleotide Database (https://www.ncbi.nlm.nih.gov/).

We computed the nucleotide and di-nucleotide frequencies in each viral genome. Among strains of the same coronavirus, some genomic sequences have long poly-A tails that are missing in other sequences. Some also have a longer 5’ untranslated region (5’ UTR) than others. To make a fair comparison between strains, the genomes were aligned with MAFFT version 7 (Katoh and Standley 2013), with the slow but accurate G-INS-1 option for 134 SARS-CoV, 20 Bovine CoV, 2 Canine CoV, 26 Murine MHV, and 10 Porcine HEV strains, and with the fast FFT-NS-2 option for large alignments for 2666 SARS-CoV-2 and 403 MERS strains. Next, using the ‘Sequence Manipulation’ tool in DAMBE7 (Xia 2018), the 5’ UTR sequences were trimmed away until the first fully conserved nucleotide position. Similarly, the 3’ UTR sequences were trimmed out up to the last fully conserved nucleotide position. Then, gaps were removed from each trimmed genome, and the global nucleotide and dinucleotide frequencies as well as their respective proportions (denoted as P_Nucleotide_ and P_Dinucleotide_, respectively) were computed in DAMBE under “Seq. Analysis|Nucelotide & di-nuc Frequency”. Additionally, nucleotide and di-nucleotide frequencies were similarly computed for whole, untrimmed, genomes. Finally, the conventional index of CpG deficiency (Cardon *et al.* 1994; Karlin *et al.* 1997) was calculated as:

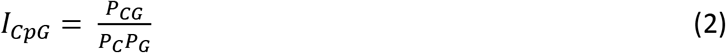

The index is expected to be 1 with no CpG deficiency or excess, smaller than 1 if CpG is deficient and greater than 1 if CpG is in excess.

Next, among 2666 high sequence quality and complete SARS-CoV-2 genomes from CNCB, we randomly selected one genome from each collection date, inclusively between December 31, 2019 (first isolate) and May 6, 2020 (most recent isolate, database last accessed on May 16, 2020), that have complete records of local region annotations and nucleotide sequences in NCBI. A total of 99 variants (or samples) were retrieved across 127 days since SARS-CoV-2 (strain Wuhan-Hu-1, MN908947) was first sequenced. For each of the 99 samples, the nucleotide sequence of 12 out of 13 viral regions (5’ UTR, ORF1ab, S, ORF3, E, M, ORF6, ORF7a, ORF8, N, ORF10, and 3’ UTR) were extracted from DAMBE in FASTA format, and local nucleotide and dinucleotide frequencies and their proportions were computed for each region. ORF7b was omitted from the analysis because it was not annotated in 30 out of 99 samples, including the reference genome Wuhan-Hu-1 (MN908947). To determine the sequence mutation patterns over time at each viral region, the nucleotide sequences from our 99 genomes were first aligned with MAFFT with the slow but accurate G-INS-1 option; then each aligned sequence was pair-wise assessed for single nucleotide polymorphisms (SNPs) using DAMBE’s “Seq. Analysis|Nucleotide substitution pattern” with reference genome = Wuhan-Hu-1 (MN908947, first sequenced) and Default genetic distance = F84.

Host tissues that are infected by SARS-CoV-2, MERS, and SARS-CoV in human, Bovine CoV in cattle, Canine CoV in dog, Murine HEV in mice, and Porcine HEV in pig were identified through an exhaustive large-scale manual search on relevant evidence-based primary source studies. The studies considered included clinical course, autopsy, and experimental infections, but cross-host studies were excluded. In total, tissue infections were determined from 25 SARS-CoV studies, 11 SARS-CoV-2 studies, 8 MERS studies, 15 Murine CoV (MHV) studies, 9 Porcine HEV studies, 18 Canine CoV studies, and 10 Bovine CoV studies (Supplemental File S1). Resultantly, the regular tissue habitats of viruses were determined based on the prevalence of virus detection in host tissues across studies. For example, among studies on SARS-CoV-2, some tissue infections (e.g., infections in the lung and intestine) are frequently recorded while others are rarely recorded (e.g., stomach). To compare the relative prevalence of SARS-CoV-2 infection, in the lung vs. in other tissues for instance, we calculated commonness of detection (COD) as:

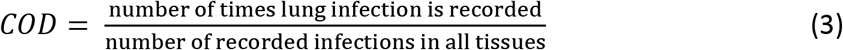

## DATA AVAILABILITY

Supplemental File S1 contains reference compilation of virus regular habitats. Supplemental File S2 and S3 contain nucleotide and di-nucleotide frequencies in trimmed and whole viral genomes, respectively. Supplemental File S4 contains the global and local mutation patterns in SARS-CoV-2 genomes. Supplemental File S5 contains the local CpG dinucleotide frequencies in a sample of 99 SARS-CoV-2 genomes. Supplemental File S6 contains Supplemental figures S1 to S5.

## RESULTS

### All host-specific coronaviruses except Murine MHV regularly infect host tissues that highly express AVPs

The tissue-specific mRNA expressions of 7 human APOBEC3 gene isoforms (A3A, A3B, A3C, A3D, A3F, A3G, and A3H) and 2 human ZAP isoforms (ZC3HAV1 and ZC3HAV1L) were retrieved from publicly available RNA-Seq datasets (see Materials and Methods) and averaged FPKM values were compared within study. Supplemental Fig. S1 shows that all 3 human coronaviruses infect the lung, heart, liver, kidney, and stomach; SARS-CoV and SARS-CoV-2 additionally infect the intestines; SARS-CoV and MERS additionally infect the skeletal muscle; but only SARS-CoV has been reported to infect the lymph node and the spleen (Supplemental Fig. S1; references with records of tissue infection are located in Supplemental File S1).

We determined which human tissues are commonly infected by coronaviruses and whether these tissues express AVPs in abundance. Figure 1 shows the commonness of detection of SARS-COV-2, SARS-CoV, and MERS (CODs) (see Equation 3, Materials and Methods) in human tissues and the relative proportions of mRNA expression (PME) of APOBEC3 and ZAP isoforms (see Equation 1, Materials and Methods) in each tissue that is susceptible to infection. Furthermore, in each susceptible tissue, the relative mRNA expressions of AVPs (in PME values) were determined as high (in green) or low (in red) (see Materials and Methods). Out of the 6 tissues with records of infection, the lung and the intestine are regular habitats of SARS-CoV-2 with the highest CODs. Furthermore, both tissues contain high PMEs for some APOBEC3 isoforms (Fig. 1a: A3A, A3B, A3D, A3G, A3H in the lung, and A3B, A3D, A3G, and A3H in the intestine) and for ZC3HAV1 (Fig. 1b). In the 4 tissues (liver, heart, kidney, and stomach) where infection is less frequently observed (less COD values), APOBEC3 and ZAP PMEs are also low (Fig. 1a, 1b). Similarly, some regular habitats of SARS-CoV-2 (lung, kidney, intestine, spleen, liver) and of MERS (lung, kidney, and liver) also host high PMEs for some APOBEC3 isoforms (Fig. 1c) and for some ZAP isoforms(Fig. 1d). Hence, all 3 human coronaviruses can regularly infect host tissues that express relatively high AVP expressions and display no strong preference for tissues with AVP deficiency.

**Fig. 1.**
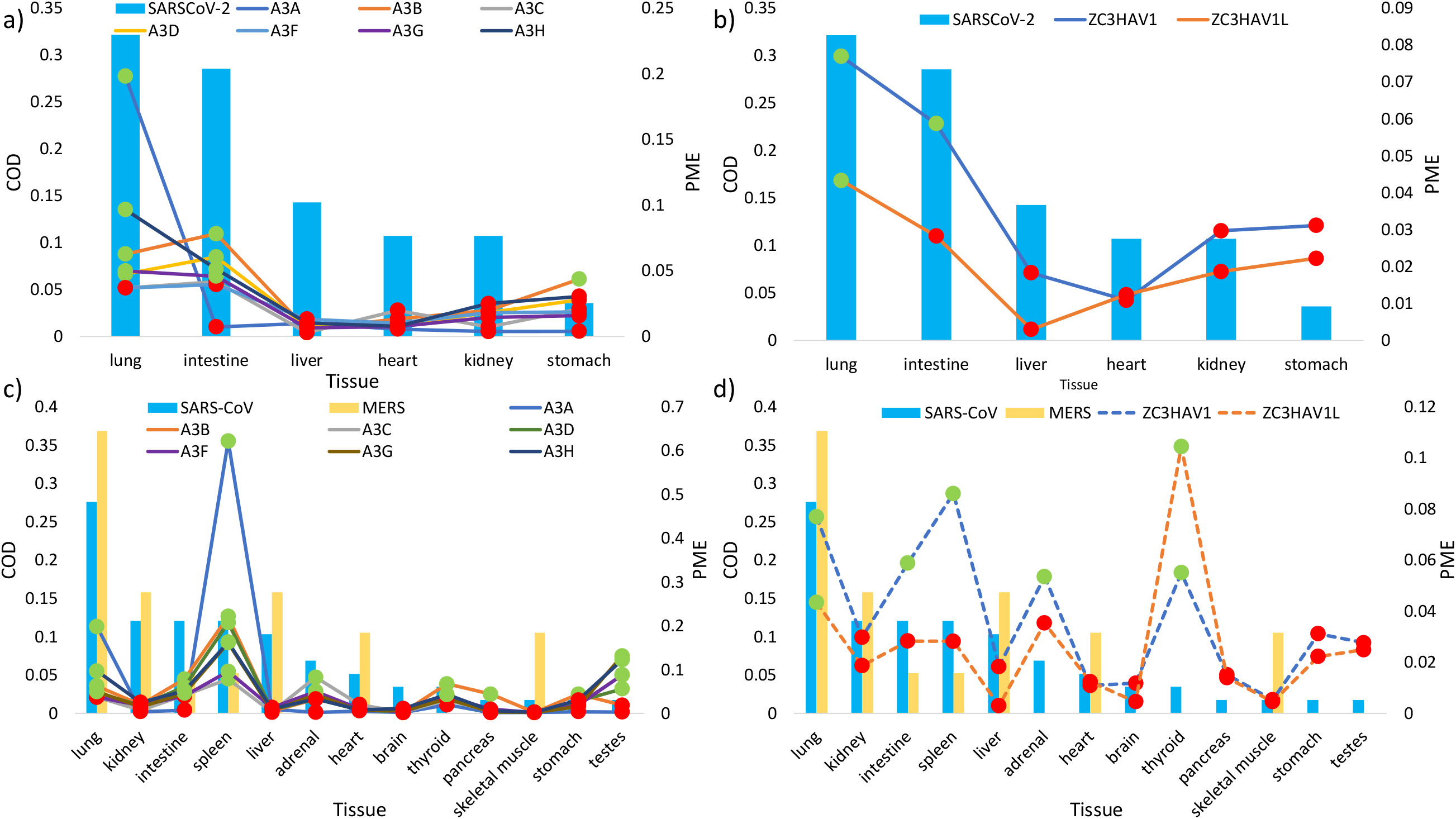
The histograms show the regular tissue habitats (as measured in commonness of detection COD, on primary Y axis) of SARS-CoV-2 (a, b) and of SARS-CoV and MERS (c, d). The lines represent the relative mRNA expression (in proportions of mRNA expression PME, on secondary Y axis) of a) APOBEC3 isoforms (solid lines) and b) ZAP isoforms (dash lines) in tissues susceptible to SARS-CoV-2 infection, and the PME of c) APOBEC3 isoforms (solid lines) and d) ZAP isoforms (dashed lines) in tissues susceptible to SARS-CoV and MERS infections. Highlighted in green and red are PME values that are greater and lower than the averaged PME values, respectively. PME values were calculated based on averaged mRNA FPKMs retrieved from the GTEx Portal (Lonsdale *et al.* 2013).

Similarly, we retrieved averaged mRNA expression levels of AVP isoforms in four other mammalian species (cattle, dog, pig, mice) and reference records of tissue-specific infections of their coronaviruses (Supplemental File S1, Supplemental Fig. S2). We determined the regular habitats (by COD) for Porcine HEV in pig (Fig. 2a), Canine CoV in dog (Fig. 2b), Bovine CoV in cattle (Fig. 2c), and Murine MHV in mice (Fig. 2d), as well as the relative mRNA expressions (PMEs) for AVP isoforms in infected tissues. Like human coronaviruses, other mammalian coronavirus can regularly infect tissues that exhibit both high APOBEC3 and ZAP mRNA expressions, such as Porcine HEV infecting pig liver (Fig. 2a), Canine CoV infecting dog intestine and lung (Fig. 2b), and Bovine CoV infecting cattle intestine (Fig. 2c). All three of these coronaviruses do not avoid tissues with high AVP expressions, nor do they display a compelling preference for tissues with low AVP expressions. Lastly, Murine MHV regularly infects mice brain and liver but rarely infects the lung; however, mice brain and liver express low levels of both APOBEC3 and ZAP PMEs, whereas the lung expresses high levels of both AVP PMEs (Fig. 2d). Hence, unlike the other coronaviruses, Murine MHV seems to avoid tissues with high AVP expressions and prefers to infect tissues with low AVP expressions.

**Fig. 2.**
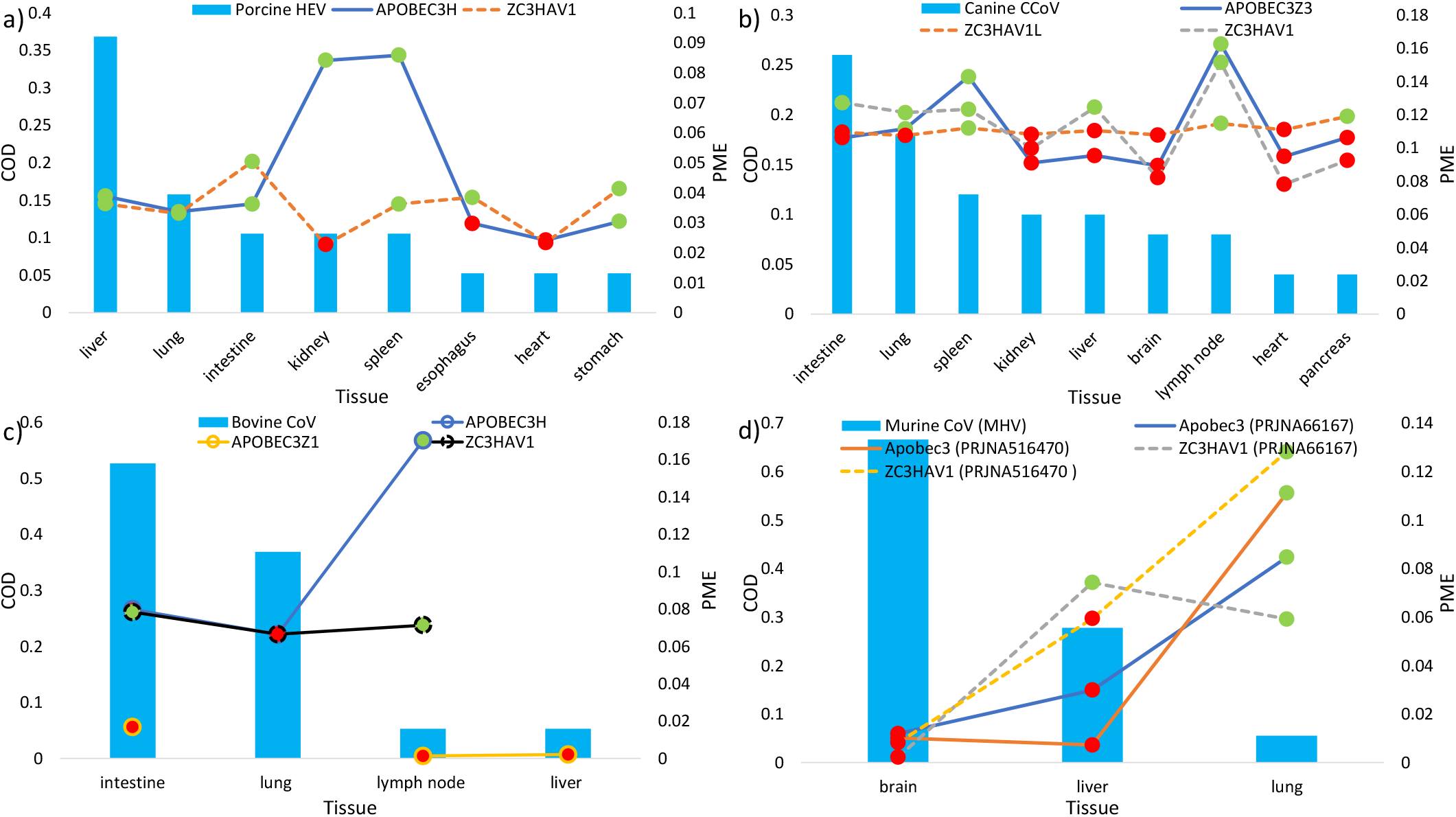
The histograms show the regular tissue habitats (as measured in commonness of detection COD, on primary Y axis) of a) Porcine HEV, b) Canine CoV, c) Bovine CoV, and d) Murine MHV. The lines represent the relative mRNA expression (in proportions of mRNA expression PME, on secondary Y axis) of (a) APOBEC3 and ZAP isoforms in pig tissues susceptible to Porcine HEV infection, PME of (b) APOBEC3 and ZAP isoforms in dog tissues susceptible to Canine CoV infections, PME of (c) APOBEC3 and ZAP isoforms in cattle tissues susceptible to Bovine CoV infection, and PME of (d) APOBEC3 and ZAP isoforms in mice tissues susceptible to Murine MHV infection. Highlighted in green and red are PME values that are greater and lower than the averaged PME values, respectively. Solid lines and dashed lines represent APOBEC3 and ZAP isoforms, respectively. PMEs were calculated based on averaged mRNA expressions retrieved from the Bovine Genome Database (Shamimuzzaman *et al.* 2019), BioProject PRJNA124245 (Briggs *et al.* 2011), TISSUE 2.0 integrated datasets (Palasca *et al.* 2018), mouse ENCODE consortium (Yue *et al.* 2014) and BioProject PRJNA516470 (Naqvi *et al.* 2019).

In principle, there are three possible classes of AVP expression a given tissue may conform to: 1) overall AVP abundance, 2) overall AVP deficiency, and 3) selective expression of AVPs. The first two classes describe tissues for which both ZAP and APOBEC3 are expressed highly and lowly, respectively. The third pattern can be divided into two subsets: one in which APOBEC3 enzymes are highly expressed and ZAP is lowly expressed, and the inverse to this pattern. Figures 1 and 2 suggest that tissue-specific APOBEC3 and ZAP expressions may be correlated in some species but not in others. Indeed, based on 24 human tissue, PME values of human APOBEC3 and ZAP are significantly positively correlated (e.g., A3H vs ZC3HAV1: R^2^ = 0.43, P = 0.00035). Similarly, we found significant positive correlation between the two AVPs in PME values based on 17 mice tissues (Apobec3 vs ZC3HAV1: R^2^ = 0.49, P = 0.0017) and based on 10 dog tissues (APOBEC3Z3 vs ZC3HAV1: R^2^ = 0.56, P = 0.021). However, there are no significant correlation between the two AVPs in PME values based on 26 cattle tissues (APOBEC3H vs ZC3HAV1: R^2^ = 0.22, P = 0.34) or based on 33 pig tissues (APOBEC3H vs ZC3HAV1: R^2^ = 0.11, P = 0.065).

### Only coronaviruses targeting tissues with high AVP expressions exhibit decreased CpG and increased U content

In the previous section, we demonstrated that many surveyed host-specific coronaviruses commonly infect tissues that exhibit high levels of AVPs (Fig. 1, 2; supplemental Fig. S1, S2), but MHV does not conform to this observation (Fig. 2d). Here we compared the CpG and U content of these coronaviruses and found that viruses that regularly infect AVP-rich tissues tend to exhibit diminished CpG content in tandem with elevated U content. Conversely, MHV neither targets AVP-rich tissues, nor does its genome indicate directional mutation with respect to CpG or U content. Both trimmed genomes (Fig. 3a, see Materials and Methods) and whole genomes (Supplemental Fig. S3a) shows that MHV has the highest I_CpG_ (about 0.6 or higher) while SARS-CoV-2 has the lowest I_CpG_ (below 0.43 in all but two genomes). As for all other coronaviruses surveyed, they also exhibit low I_CpG_ < 0.5 except for MERS being slightly higher. It should be noted that among the 7 coronaviruses surveyed, I_CpG_ values also show the greatest variation among Murine MHV genomes whereas I_CpG_ values are relatively much more constrained among genomes of the other 6 (Supplemental Fig. S4a). Indeed, CpG content is weakly constrained in Murine MHV.

**Fig. 3.**
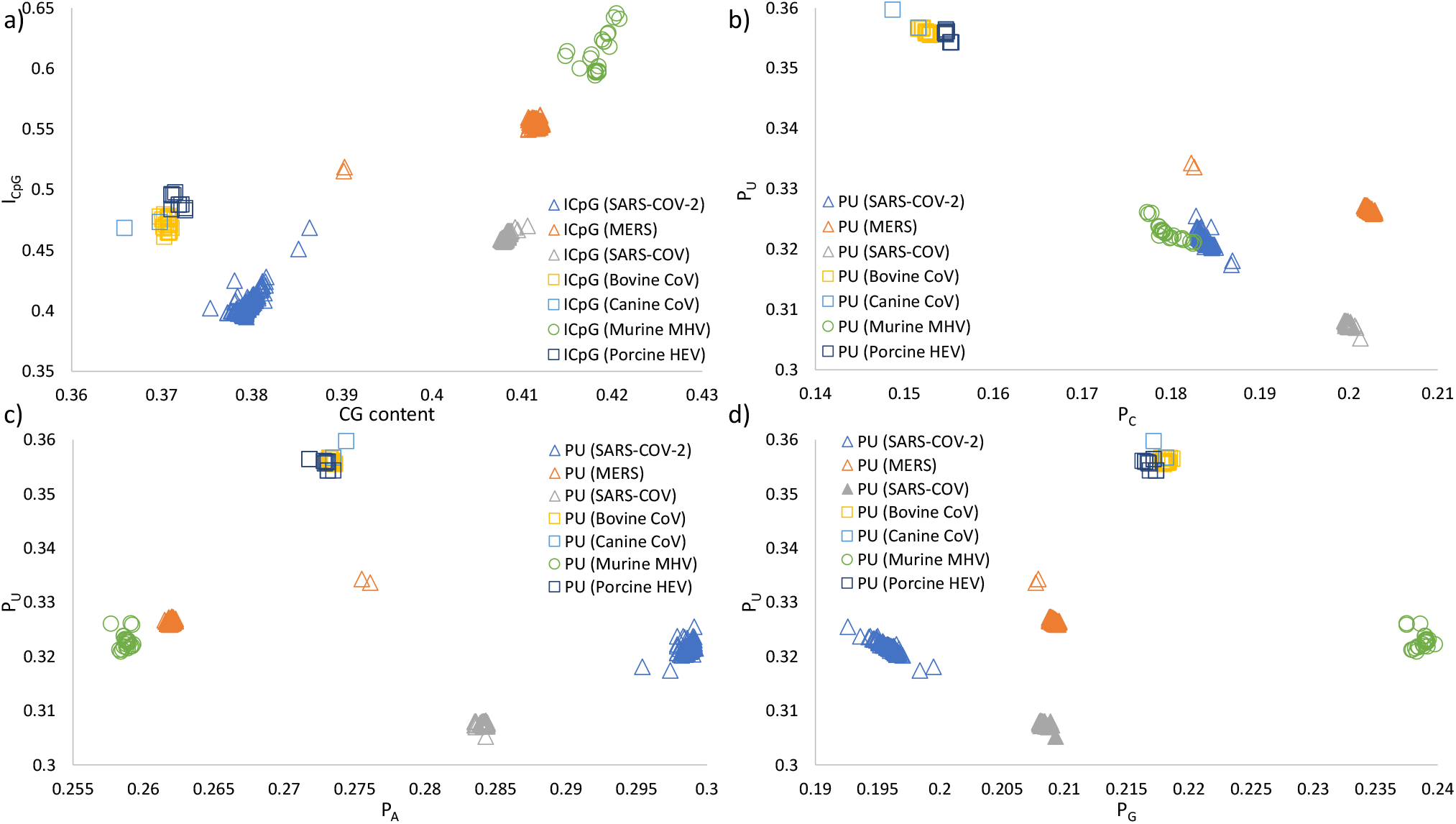
ICpG and nucleotide proportions for seven Coronaviruses with complete genomes and host information. All genomes were aligned with MAFFT and sequence ends were trimmed (see Materials and Methods). In panel a) shows that SARS-CoV-2 has the least ICpG in comparison to other coronaviruses from their natural hosts. In panels b), c) and d) show that proportions of U (P_U_) negatively correlates with PC but not with P_A_ or P_G_, and that P_U_ is similarly the highest among Bovine CoV, Canine CoV, and Porcine CoV, and similarly the lowest among Murine MHV and human coronaviruses. Each panel includes 2666 SARS-CoV-2 genomes, 403 MERS genomes, 134 SARS-CoV genomes, 20 Bovine CoV genomes, two Canine CoV genomes, 26 Murine MHV genomes, and 10 Porcine HEV genomes.

Figure panels 3b, 3c, and 3d show that the proportion of U nucleotides (P_U_) decreases with the proportion of C nucleotides (P_C_), but P_U_ does not correlate with P_A_ or P_G_. This global relationship in the trimmed genomes may suggest a hallmark of C to U deamination in coronaviruses, that single stranded RNA genomes could indeed be subjected to editing by APOBEC3 proteins. Specifically, Bovine CoV, Canine CoV, and Porcine HEV all have very high P_U_ and conversely very low P_C_. In comparison, since Murine MHV does not infect host tissues with high APOBEC3 expression, it may have been subjected to less C to U deamination and therefore it has notably reduced P_U_ and increased P_C_. Additionally, similar to I_CpG_, P_U_ is least constrained in Murine MHV in comparison to any other coronavirus (Supplemental Fig. S4b). As for human coronaviruses, figure 3b shows that P_U_ is comparably low in SARS-CoV-2 and in MERS, like in Murine MHV, and even lower in SARS-CoV. Nonetheless, it is important to note that the emergence of all three human coronaviruses are much more recent in comparison to coronaviruses of other mammals; their genomes had short evolutionary time to be shaped by host AVPs. Lastly, the same patterns were observed when P_U_ was re-analyzed using whole, untrimmed, genomes (Supplemental Fig. S3).

### Strong evidence of directional mutation shaped by AVPs in local SARS-CoV-2 viral regions

In the above results, we demonstrated that viral genomes show more pronounced shifts towards CpG deficiency and elevated U content when the virus regularly infect host tissues with high expression of the two AVPs. However, two limitations of figure 3 are 1) it does not show the substitution patterns within local viral regions (such as the ORF1ab region), and 2) because human viruses share similar or lower global P_U_ in comparison to Murine MHV which predominantly infects AVP-deficient tissues, we cannot suggest that APOBEC3 would shape human coronavirus genomes over time. To resolve these limitations, we examined whether there has been an evolutionary history of P_U_ elevation in local SARS-CoV-2 regions over the span of 4 months since the virus was first isolated.

To resolve the first limitation, figure 4 shows single nucleotide polymorphisms (SNPs) between 28475 aligned SARS-CoV-2 sequences (including complete and incomplete sequences) (retrieved from https://bigd.big.ac.cn/ncov/variation/statistics, last accessed May 16, 2020), and the comparison was made against the first identified Wuhan-Hu-1 strain (MN908947). Based on global sequence comparison, figure 4a shows that most SNPs are C->U substitutions. More specifically, local mutation patterns (Fig. 4b) show that among 28475 sequence samples, C->U substitutions are most prevalent at the 5’ UTR region and the ORF1ab region (normalized by region length), but not at any other viral regions. To resolve the second limitation, figure 5 and supplemental figure S5 show the local mutation patterns over time in a sample of 99 complete and high-quality SARS-CoV-2 genomes with complete NCBI annotations. Each retrieved sample had been collected on a different day, since first isolation (Wuhan-Hu-1, MN908947, December 31, 2019) to the most recently isolated sample (mink/NED/NB04, MT457401, May 6, 2020) (see Materials and Methods), and each sample was grouped into one of six time ranges. Indeed, with reference to strain Wuhan-Hu-1, aligned sequences show an excessive number of C->U substitutions, and that the total number of C->U substitutions increases over time, but only at the 5’ UTR region (Fig. 5a) and ORF1ab region (Fig. 5b) and not in other regions (Fig. 5c, 5d, Supplemental Fig. S5). It is noteworthy that in the S region, directional mutation over time favours A->G substitutions; whereas in the ORF3a region, directional mutation over time seems to favour G->U, but the number of G->U mutations have decreased in the latest group of genome samples. Together, figures 4 and 5 suggest that APOBEC3 may indeed edit single-stranded RNA genomes, specifically the 5’UTR and ORF1ab regions in SARS-CoV-2. Whereas in the S region specifically, A->G directional mutation may be the result of deamination by the mammalian adenosine deaminase acting on RNA type 1 (ADAR1) enzyme (Jiang 2020; Samuel 2011; Zhao *et al.* 2004).

**Fig. 4.**
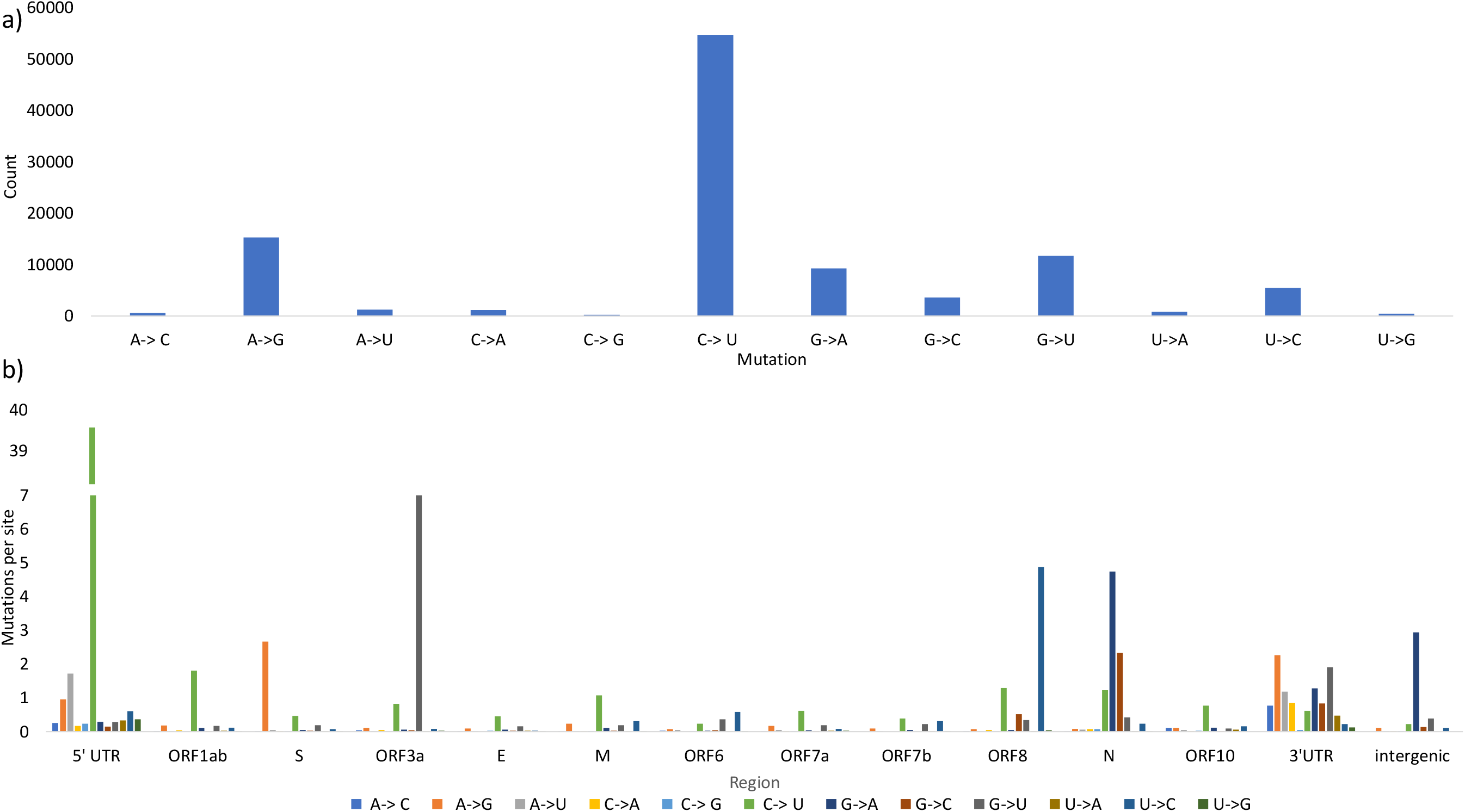
Single nucleotide polymorphisms in 28474 SARS-CoV-2 (complete and incomplete) samples sequenced to date (May 6, 2020), with reference to strain Wuhan-Hu-1 (MN908947). Panel a) shows strong global C->U directional mutation when the entire genomes are considered. Panel b) shows that the number of C->U directional mutations (normalized by the length of the region) is predominantly observed in the 5’ UTR and ORF1ab viral regions but not in other regions. Indels and ambiguous point mutations were omitted.

**Fig. 5.**
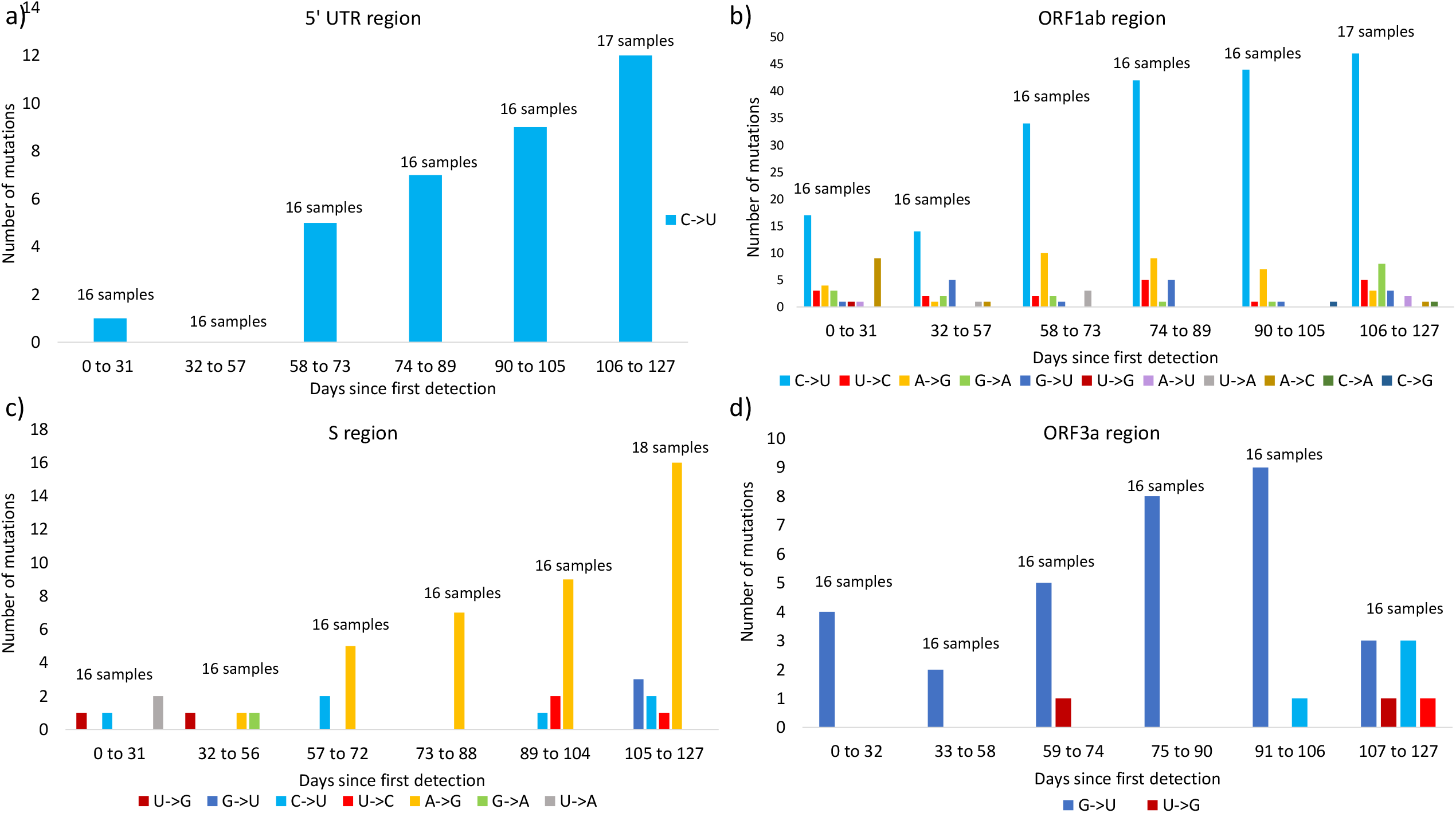
Local mutation patterns over time in a sample of 99 complete and high-quality SARS-CoV-2 sequences with complete NCBI annotations. Each retrieved sample was collected on a different day, since first isolation (Wuhan-Hu-1, MN908947, December 31, 2019) to the most recent isolation (mink/NED/NB04, MT457401, May 6, 2020) (see Materials and Methods), then each sample was grouped into one of six time ranges. In panels a) and b) shows that the total number of C->U mutations are the most prevalent and they increase over time, in the 5’ UTR region and ORF1ab region, respectively. In panels c) and d) show that C->U mutation are not prevalent, and that A->G mutation and G->U mutations are favoured in the S and ORF3a regions, respectively.

Lastly, we compared differences in I_CpG_ between viral regions over time among the 99 SARS-CoV-2 samples. Figure 6 shows no difference in I_CpG_ between sequences sampled at different time (Since December 31, 2019 to May 6, 2020). This suggests that I_CpG_ changes slowly over time. However, there are notable differences in ICpG among specific viral regions. In particular, ORF1ab, S, and ORF6 regions have the lowest I_CpG_ values, whereas the 5’ UTR, E, and ORF10 regions have the highest ICpG values at above 1. The selective pressure for CpG deficiency to evade ZAP is not uniform across different SARS-CoV-2 viral regions.

**Fig. 6.**
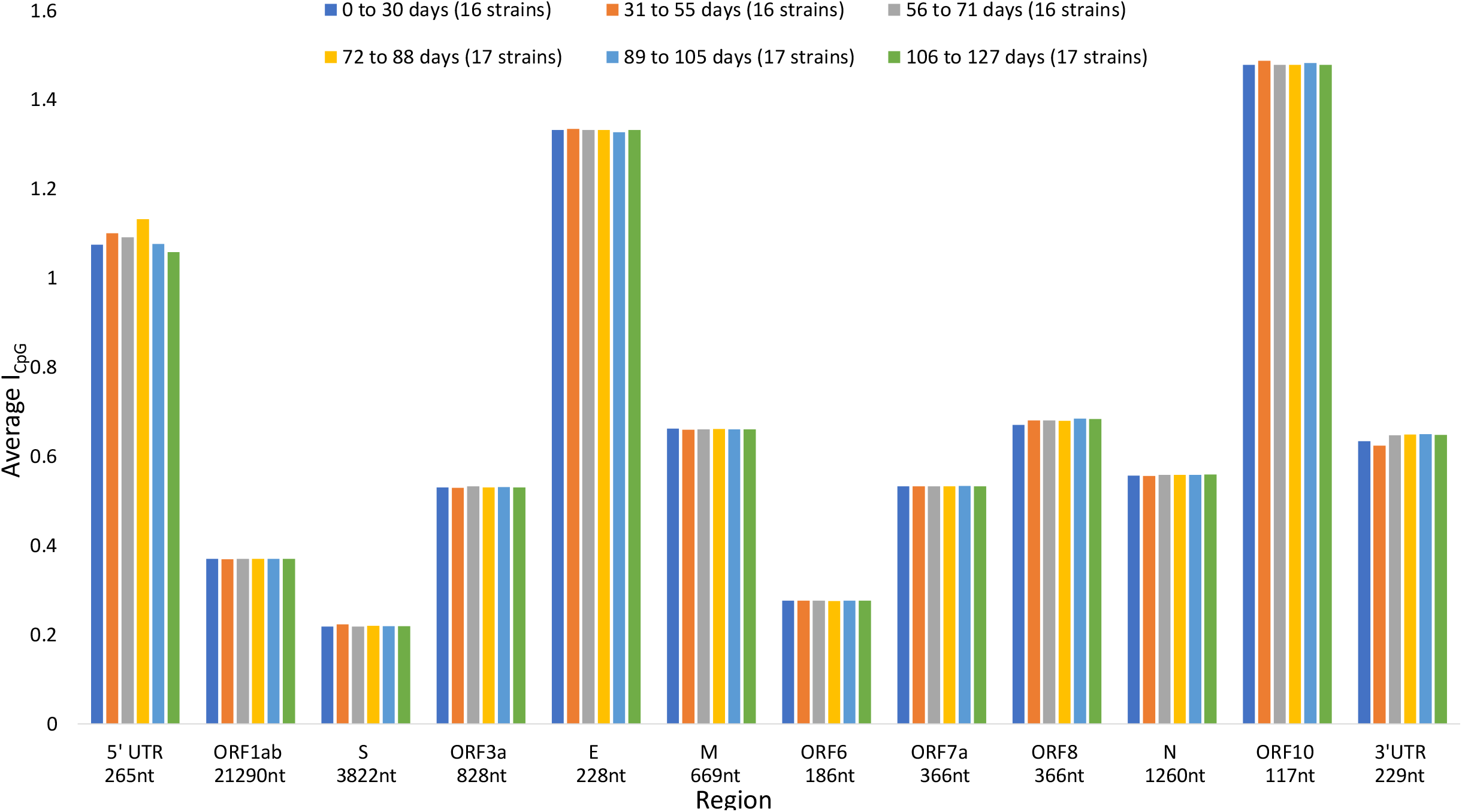
Local I_CpG_ values over time in a sample of 99 complete and high-quality SARS-CoV-2 sequences with complete NCBI annotations. Each retrieved sample was collected on a different day, since first isolation (Wuhan-Hu-1, MN908947, December 31, 2019) to the most recent isolation (mink/NED/NB04, MT457401, May 6, 2020) (see Materials and Methods), then each sample was grouped into one of six time ranges. I_CpG_ does not change substantially over the 127 days since first detection, but I_CpG_ is not uniform across viral regions. I_CpG_ is the lowest in ORF1ab, S, and ORF6 regions, and the highest in the 5’ UTR, E, and ORF10 regions.

## DISCUSSION

SARS-CoV-2 poses a serious global health emergency. Since its outset in Wuhan City, Hubei province of China in December 2019, the viral outbreak has resulted in over 7 million confirmed cases around the world (https://www.who.int/emergencies/diseases/novel-coronavirus-2019, last accessed June 11, 2020). The pandemic has prompted an immediate global effort to sequence the genome of SARS-CoV-2, and over 28000 genome samples have been publicly deposited over the course of just four months to facilitate vaccine development strategies. With a wealth of sequence data, we performed a comprehensive comparative genome study on SARS-CoV-2 and six other coronaviruses across five mammalian species, with the aim to understand how coronaviruses evolve in response to tissue-specific host immune systems.

We tested the hypothesis that both APOBEC3 and ZAP immune responses act as primary selective pressures to shape the genome of an infecting coronavirus over the course of its evolutionary history within host tissues. Specifically, viral genomes are driven towards reduced CpG dinucleotides to elude ZAP-mediated cellular antiviral defense, and increased U residues because of RNA editing by APOBEC3 proteins. In line with our expectations, we found compelling hallmarks of CpG deficiency and C to U deamination globally in mammalian coronaviruses (i.e., Fig. 3: Bovine CoV, Canine CoV, and Porcine HEV) that regularly infect host tissues expressing both AVPs in abundance (Fig. 2a, 2b, and 2c). Unsurprisingly, these global trends were absent from Murine MHV genomes (Fig. 3) as this virus does not regularly infect tissues that highly express AVPs (Fig. 2d). Corroborating this observation, both I_CpG_ and P_U_ values show the greatest variation among Murine MHV genomes (Supplemental Fig. S4), suggesting that MHV is not functionally constrained by either AVP. This aligns with our prediction that for a virus regularly infecting host tissues that are deficient in AVPs, there will be no strong directional mutations resulting in decreased CpG dinucleotides or elevated U residues. Conversely, when a virus regularly infects host tissues that are abundant in ZAP and APOBEC3, these AVPs shape the molecular evolution of viral genomes in two ways: CpG deficiency contributes to the survival of the virus by evading ZAP-mediated antiviral defense through CG dinucleotide recognition, and elevated U content as the result of genome editing by APOBEC3.

In comparison to other mammalian coronaviruses, human coronaviruses (SARS-CoV-2, SARS-COV, MERS) have been circulating in the human hosts for a much shorter time, particularly SARS-CoV-2. Among the three, SARS-CoV-2 genomes shows extreme CpG deficiency (Fig. 3a); its I_CpG_ values are comparable to that of the Bat CoV RaTG13 coronavirus infecting the bat species *Rhinolophus affinis* (Xia 2020) but lower than that of all other coronaviruses studied herein as well as all other mammalian specific coronaviruses (Xia 2020). Indeed, many recently published studies point to Bat CoV RaTG13 as the most closely related known relative of SARS-CoV-2 when the whole genome is considered (Andersen *et al.* 2020; Lai *et al.* 2020; Shang *et al.* 2020; Tang *et al.* 2020), and to *Rhinolophus affinis* as a potential intermediate host or reservoir for SARS-CoV-2 (Liu *et al.* 2020). Moreover, local comparative analyses on CpG content have two implications. First, SARS-CoV-2 has acquired CpG deficiency in an intermediate reservoir prior to zoonotic transmission to humans, as CpG deficiency may be acquired slowly since there is no notable change in I_CpG_ across all 12 viral regions in the span of four months since SARS-CoV-2 was initially isolated. In this context, it is regrettable that the Bat CoV RaTG13 was not sequenced when it was initially sampled in 2013. The downshifting in I_CpG_ in RaTG13 would have served as a warning that the virus will likely infect tissues with high ZAP expression, because the viral genome has successfully evolved to evade ZAP-mediated antiviral defense in humans. Second, the evolutionary pressure for CpG deficiency may be region specific. The S, ORF1ab, and ORF6 regions have the most severe CpG deficiencies (Fig. 6, I_CpG_ < 0.4), whereas the 5’ UTR, E, and ORF10 regions have the highest CpG content with no signs of CpG deficiency (Fig. 6, I_CpG_ > 1). While evolution has allowed the Spike protein to elude ZAP because it is crucial for host cell recognition and entry, structural genes such as the Envelope and Membrane protein are subjected to less selective pressure to evade ZAP.

A current survey of SARS-CoV-2 genomes does not indicate drastically increased U and decreased C contents. A global sequence comparison shows that SARS-CoV-2 (and SARS-CoV and MERS) have comparable U and C contents as Murine MHV, but higher U and lower C contents in comparison to Bovine CoV, Canine CoV, and Porcine HEV (Fig. 3b). This is because while a coronavirus infecting a specific host tissue for a long time would experience the same cellular antiviral environment and is consequently expected to have undergone significant RNA editing; newly emerging coronaviruses such as SARS-CoV-2 would not have enough time to accumulate a high number of RNA modifications. Nevertheless, global nucleotide substitution patterns (Fig. 4a) show that C to U substitution is still the most prevalent among SARS-CoV-2 genomes collected to date. This prevalence in genome wide C to U substitutions in SARS-CoV-2 has been similarly reported by Di Giorgio *et al.* (2020), who also observed the same trends in SARS-CoV and to a lesser degree in MERS.

More importantly, local sequence comparisons among SARS-CoV-2 samples indeed show that there is an evolutionary history of P_U_ elevation in specific SARS-CoV-2 viral regions over the span of 4 months since the virus was first isolated. There is an excessive number of C to U substitutions, and the prevalence of C to U mutations is increasing over time, specifically in the 5’ UTR and ORF1ab regions (Fig. 4b, 5a, 5b). This implies that these two specific viral regions are under constant C to U deamination by the APOBEC3 gene family, at least in the short term so far. Another noteworthy observation is that G to A substitution is preferred in the S region and the numbers of G to A substitutions are increasing over time (Fig. 5c). The preference for this mutation may be caused by deamination by the mammalian adenosine deaminase acting on RNA type 1 (ADAR1) enzyme (Di Giorgio *et al.* 2020; Jiang 2020), which edits A into I, and subsequently into G. Although, ADAR1 was known for targeting double-stranded RNAs, not single-stranded RNA sequences (Eisenberg and Levanon 2018; O’Connell *et al.* 2015; Simmonds 2020; Zhao *et al.* 2004). Regardless, these results suggest that RNA editing by host deaminase systems may indeed act on coronaviruses.

While it is important to determine the evolution of coronavirus genomes to understand its host adaptation and specificity, this study focuses more on the evolutionary pressure and RNA editing process that host immune systems exert onto viral genomes. Our aim is to prompt motivations for vaccine designs in the development of attenuated pathogenic RNA viruses. Previous experimental works have shown that increasing CpG dinucleotides in CpG-deficient viral genomes leads to drastic decrease in viral replication and virulence (Antzin-Anduetza *et al.* 2017; Burns *et al.* 2009; Fros *et al.* 2017; Trus *et al.* 2020; Tulloch *et al.* 2014; Wasson *et al.* 2017), and in recent years several studies have proposed vaccine development strategies involving increased CpG to attenuate pathogenic RNA viruses (Burns *et al.* 2009; Ficarelli *et al.* 2020; Trus *et al.* 2020; Tulloch *et al.* 2014). Among coronaviruses, SARS-CoV-2 has the most extreme CpG deficiency (Xia 2020), particularly in the S protein coding region (Fig. 6). Increasing CpG content at the S protein may provide a good starting point for strategies to inhibit SARS-CoV-2’s ability to recognize and enter host cells. On the other hand, because C to U deamination cannot be proof-read by viral exonuclease Nsp14-ExoN (Eckerle *et al.* 2010; Smith *et al.* 2013; Victorovich *et al.* 2020), host innate deaminases may drive up the rate of evolution in viral genomes (Di Giorgio *et al.* 2020) or modify CpG into UpG to further increase CpG deficiency and reduce viral susceptibility by ZAP. The possibility of APOBEC3 editing activity acting on RNA viruses and its potential exploits by viruses such as SARS-CoV-2 in the long term require further investigation and scrutiny.

## ACKNOWLEDGEMENTS

This work is supported by the Natural Sciences and Engineering Research Council of Canada (NSERC) Discovery Grant to X.X. [RGPIN/2018-03878], and NSERC Doctoral Scholarship to Y.W. [CGSD/2019-535291].

## AUTHOR CONTRIBUTIONS

Y.W. and X.X. designed the study. Y.W., J.R.S., and X.X. wrote the manuscript. Y.W., P.A. and X.X. collected the data. Y.W. and J. R. S. analyzed the data. Y.W., P.A., and J. R. S. prepared all figures. All authors reviewed the manuscript. X.X. supervised the project.

